# Aortic Valve Remodeling in Chronic Kidney Disease: A Mineralocorticoid Receptor–Driven Mechanism Involving NGAL and TLR4 Pathways

**DOI:** 10.1101/2025.09.12.675970

**Authors:** T. Sánchez-Bayuela, M. Soulié, M. Garaikoetxea, A Fernández-Celis, N. LópezAndrés, F Jaisser

## Abstract

**INTRODUCTION:** Aortic stenosis (AS) is the most prevalent valve heart disease. Renal failure increases the risk of AS and many circulating factors released during chronic kidney disease (CKD) participate to AS pathophysiology. We assessed in this study the role of increased aldosterone levels occurring in CKD as well as MR signaling and its interplay with Neutrophil Gelatinase-Associated Lipocalin (NGAL) in aortic valve interstitial cells (VICs) pathophysiology.

**METHODS:** We conducted *in vivo* studies using a CKD rat model, including both wild-type (WT) and NGAL knockout (KO-NGAL) animals, for subsequent *ex vivo* analysis of aortic valves. In parallel, primary rat VICs were used *in vitro* to assess osteogenic, fibrotic, and inflammatory responses to aldosterone, as well as to identify the key signaling pathways involved. qPCR, Western blot, and ELISA were employed to characterize these pathways.

**RESULTS:** Our findings demonstrate that MR signaling plays a central role in AS progression during CKD, as well as in VIC differentiation, inflammation, fibrosis, and calcification, mediated via the TLR4–MyD88 innate immunity pathway in aldosterone-induced responses. Furthermore, NGAL was shown to act downstream of MR to activate TLR4, promoting additional remodeling and calcification in aortic valves and VICs. These results were further validated in human samples from CKD patients.

**CONCLUSION:** Overall, this study identifies a novel signaling pathway in AS pathophysiology in the CKD rat model and *in vitro* systems, highlighting for the first time the interplay between MR, NGAL, and TLR4 in driving the pathological processes underlying AS in CKD.

**GRAPHICAL ABSTRACT:** https://BioRender.com/j7vufds

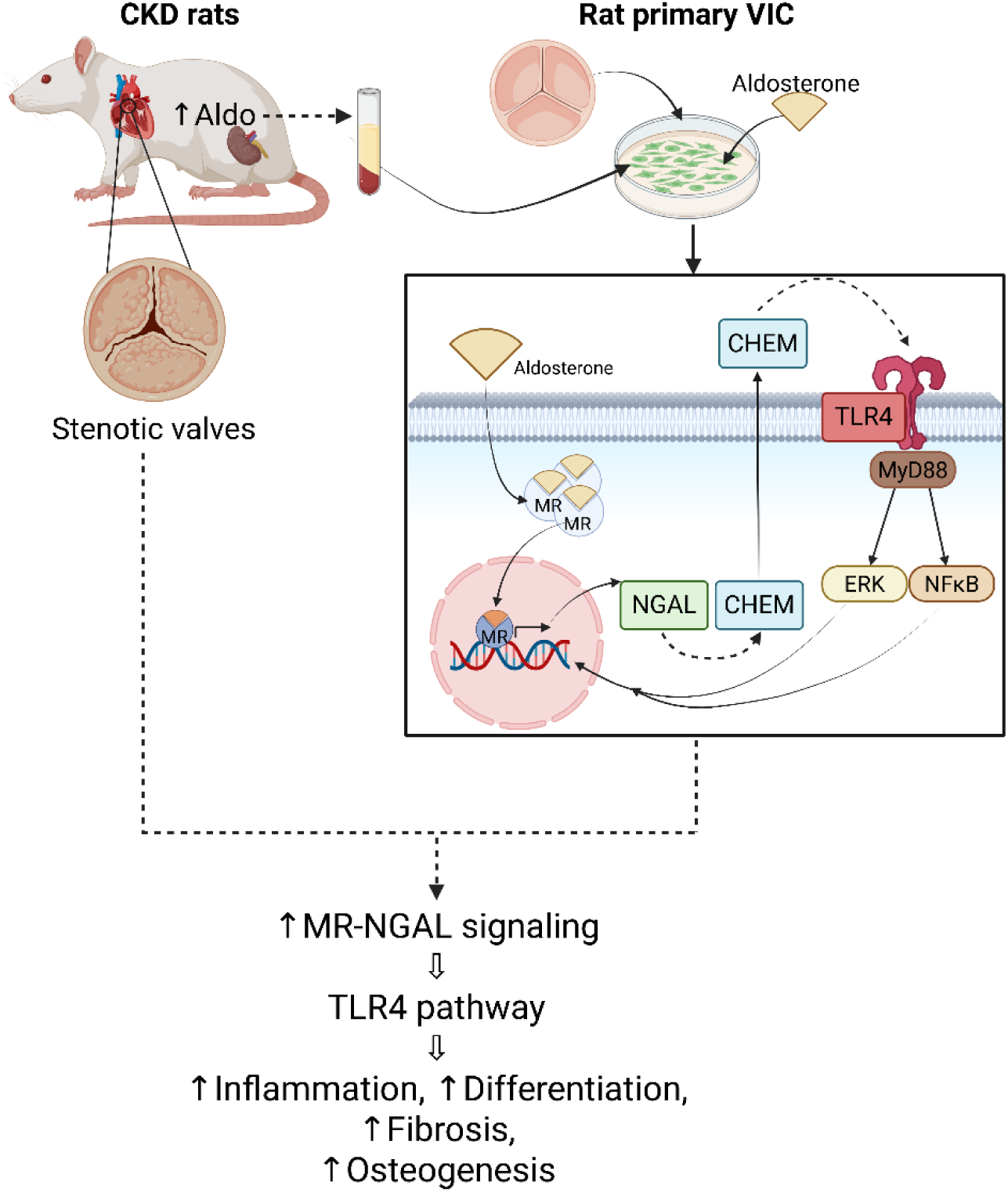

## INTRODUCTION

Aortic stenosis (AS) is the most prevalent valve heart disease in the elderly population of developed countries, with a projected disease burden expected to increase from 2.5 million in 2000 to 4.5 million in 2030^1^. The only treatments currently available are surgical valve replacement and transcatheter aortic valve (AV) implantation^1^.

The initial stages of AS resemble those of atherosclerosis; (i) endothelial disruption, (ii) renin-angiotensin system activation, (iii) lipid deposition and (iv) inflammation, progressing from immune cell infiltration to cytokine secretion and the release of pathogen-associated molecular patterns. Several inflammatory pathways play an important role in the initial stages of AS. The toll-like receptors (TLRs) of innate immunity, which signal through the NFκB and mitogen-activated protein kinase (MAPK) pathways, regulate the expression of some pro-inflammatory and osteogenic genes^2,3^, leading to valve cell differentiation, fibrosis, osteogenesis and microcalcification. Later stages of AS involve structural changes and macrocalcification^4^.

The risk factors associated with AS include hypertension, advanced age, male sex, high cholesterol levels, smoking, renal failure, diabetes, and lipid accumulation such as high levels of low-density lipoproteins (LDL) or lipoprotein (a)^4^. Chronic kidney disease (CKD) is characterized by a gradual decline in kidney function, leading to a significant decrease in estimated glomerular filtration rate and an increase in blood pressure in the long term. CKD and AS are interrelated, each affecting the progression and prognosis of the other. An estimated 10-15% of individuals with CKD develop AS, whereas 30-35% of individuals with AS also suffer from CKD or renal failure^5^. Indeed, AV calcification is highly prevalent in CKD patients and appears to progress more rapidly in patients with end-stage renal disease than those with normal renal function, with an inverse correlation between the rate of stenosis progression and glomerular filtration rate^6^.

Chronic kidney disease (CKD) is characterized by elevated circulating levels of aldosterone (Aldo) due to dysregulation of the renin–angiotensin system, a condition positively correlated with disease progression^7,8^. Activation of the mineralocorticoid receptor (MR) by aldosterone has been linked to renal and vascular fibrosis, inflammation, and calcification^9,10^. MR expression in rabbit and human AVs has been linked to the development of aortic stenosis (AS)^11,12^. Additionally, advanced stages of CKD are associated with increased concentrations of calcium and phosphate ions and complexes, as well as uremic toxins, all of which may adversely affect AV structure and function^7,13^.

Neutrophil gelatinase-associated lipocalin (NGAL) is a small, secreted protein involved in the immune response, iron transport, and cell differentiation. It is a well-established biomarker of CKD progression and is directly involved in renal inflammation and fibrosis via aldosterone/MR activation^14,15^. It is also upregulated in the AV during early stages of AS, particularly under hypoxic conditions^16^. Moreover, NGAL has been identified as a downstream effector of MR signaling in the heart during myocardial infarction and in the AV of diabetic men ^17,18^. In line with this, VICs have been described to be a source of NGAL in AS^18^. Of interest, NGAL overexpression is associated with enhanced inflammatory, oxidative stress, osteogenic and calcific burden in AS^18^. In this context, chemerin (*RARRES2*), a recently identified downstream target of MR, has been found to mediate valvular interstitial cell (VIC) differentiation, inflammation, and calcification through the MR–NGAL signaling axis in the AVs of diabetic men^19^.

These findings suggest that aldosterone/MR activation promotes the progression of AS associated with CKD, and that MR downstream effectors, such as NGAL and chemerin, are key contributors to the pathological crosstalk between inflammation, fibrosis, and calcification in the aortic valve.

The aim of this study was, therefore, to determine the role of MR activation and to identify the signaling cascade responsible for mediating the interplay between CKD and AV during AS pathophysiology.

## MATERIALS AND METHODS

### Patient population

Studies involving human valves were approved by the competent research ethics committee (Pyto.2013/26, no. 137) in accordance with Spanish legislation (BPCCPMP/ICH/135/95) and the ethical principles outlined in the Declaration of Helsinki (1975) and its amendments. All participants provided written informed consent. Human male AVs were obtained from patients undergoing elective surgical valve replacement at the *Hospital Universitario de Navarra* (Pamplona, Spain). The criteria for inclusion in the study are detailed in the **Supplementary Methods**.

### Animals

Experiments were approved by the Darwin ethics committee of Sorbonne University (#22206–2019110515362968 v4) and conducted in accordance with INSERM animal care rules and DIRECTIVE 2010/63/EU of the European Parliament. Animals were housed in a controlled-climate facility with a 12-h light/12-h dark cycle and free access to food and water.

CKD was induced by subtotal nephrectomy in male 12-week-old Sprague–Dawley rats obtained from Charles River, or in rats with a knockout of the NGAL gene (KO-NGAL). The *Ngal* mutant rat line was generated at the Institut Clinique de la Souris– PHENOMIN-ICS (http://www.phenomin.fr). Details are provided in **Supplementary Methods.**

### Cell isolation and culture

Hearts were collected from male rats and AVs were isolated under sterile conditions. AVs were enzymatically digested by incubation with collagenase II solution (240 U/mg tissue, Worthington Biochemical Corporation) for 1 h at 37 °C. Valve interstitial cells (VICs) were pelleted at 350 x *g* for 5 minutes and were then expanded in DMEM/F12 (Lonza) supplemented with 20% FBS (Gibco), 100 U/mL penicillin (Lonza), 100 µg/mL streptomycin (Lonza), 10 ng/mL FGF-2 (R&D) and 5 μg/mL insulin (Sigma-Aldrich) in 2% gelatin-coated flasks. Cells were cultured at 37 °C in an incubator, under an atmosphere containing 5% CO_2_, and were used between passages 3 and 6.

Human male VICs were isolated and expanded from AVs harvested during elective surgical valve replacement interventions (6 biological replicates/sex), as previously published^20^. In brief, AV were minced and incubated in a Collagenase type 2 (240 U/mg) buffered-solution (Worthington Biochemical Product) for 1 h, 37 °C in a saturation humidified 5% CO_2_ incubator. VICs were pelleted at 350 g, 5 min and expanded onto 2% gelatin-coated flasks using DMEM/F12 media (Lonza,) supplemented with 20% FBS (Gibco), 100 U/mL Penicillin (Lonza), 100 ug/mL streptomycin (Lonza), 10 ng/mL FGF-2 (Novus Biologicals) and 5 µg/mL insulin (Sigma-Aldrich). All experiments were conducted in technical triplicates unless otherwise indicated.

For experiments, human and rat VICs were grown overnight in low-charcoal-stripped serum (1%) and low-glucose (1.1 g/L) DMEM (Gibco). VICs were treated for the times indicated with aldosterone (Aldo 10^-8^ M, Sigma Aldrich), rat recombinant (r)NGAL (500 ng/mL, R&D), rat recombinant (r)chemerin (50 nM, abbexa) or pooled serum (10%) obtained from the *in vivo* experiments with sham-operated or CKD rats, generated as described above. Pharmacological MR blockade was performed by treating VICs with finerenone (Fine, 10^-6^M, Selleckchem). TLR4 inhibition was performed with resatorvid (TAK-242, 1 µM, MedChem). Chemokine-like receptor 1 (CMKLR1) was blunted by an specific inhibitor, 2-α-naphthoyl)ethyltrimethylammonium iodide (α-NETA, 10 μM, MedChem).

### Enzyme-linked immunosorbent assays (ELISA)

Levels of COL1, chemerin, IL-6, NGAL, MCP-1/CCL2, and OPN in cell culture supernatants were quantified at multiple time points following VIC stimulation, according to the manufacturer’s protocols (Table S2). Protein concentrations in human aortic valve samples were assessed using specific ELISA kits for human NGAL and chemerin (Table S2). Absorbance was measured using a Tecan Spark 10M multimode plate reader.

### RNA isolation and RTq-PCR

Total RNA was isolated with Tri reagent® (Sigma-Aldrich), and subjected to reverse transcription to generate cDNA, with an M-MLV reverse transcriptase (Thermo Fisher Scientific). Quantitative PCR analyses were then performed with Sybr iQ syber® (Bio-Rad) in a CFX Connect Real-Time PCR System. The relative expression of each selected gene product was calculated by the 2^−Δ*Ct*^ method, with *Gapdh* as a housekeeping gene. The primers used are listed in Table S3.

### Protein isolation and western blotting

Whole-cell protein lysates were obtained in cOmplete™ Lysis Buffer supplemented with cOmplete™ ULTRA protease inhibitor tablets and PhosSTOP™ phosphatase inhibitors (Roche), according to the manufacturer’s instructions. We subjected 25 µg protein per sample to denaturing SDS-PAGE on 4–15% TGX™ precast gradient gels (Bio-Rad) and transferred the resulting bands onto Hybond-C Extra nitrocellulose membranes with 0.2 µm pores (Bio-Rad). Membranes were incubated overnight at 4 °C with primary antibodies against TLR4, MyD88, phospho-NFκB, and phospho-ERK at the concentrations specified in Table S4. The densitometric quantification of immunoreactive bands was performed with Image Lab software (Bio-Rad) and signal intensities were normalized against total protein content per gel, as determined from an unstained gel run in parallel.

### Histology and immunohistochemistry

Rat AVs were embedded in paraffin and cut into 5 μm-thick sections, which were then deparaffinized and rehydrated in distilled water. Immunohistochemistry was performed with the automated Leica BOND-Polymer Refine Detection System (Leica), according to the manufacturer’s protocol. The primary antibodies and their working concentrations are listed in Table S5. Primary antibody binding was detected with poly-HRP-conjugated anti-mouse or anti-rabbit IgG secondary antibodies. Immunoreactivity was visualized with an enhanced 3,3′-diaminobenzidine (DAB) chromogenic substrate (Leica). Slides were mounted in DPX mounting medium (Merck/Sigma-Aldrich). Image quantification was performed using ImageJ software. For each protein, quantification was done in all three leaflets of the same valve, and the mean value was used to represent each AV from a different animal. IHQ data was expressed as % of positive DAB staining normalized to the total area of each leaflet

### Statistical analyses

Statistical analyses were performed with GraphPad Prism 9.0.2 for Windows (GraphPad Software, Boston, MA, USA). The normality of the data distribution was assessed with Shapiro– Wilk and Kolmogorov–Smirnov tests. Comparisons between two groups were performed with unpaired Student’s *t*-tests or Mann–Whitney *U* tests, depending on the data distribution (normal or non-normal, respectively). Comparisons between multiple groups were performed by one-way ANOVA for normally distributed data or with Kruskal–Wallis tests for data that were non-normally distributed (even after transformation), followed by appropriate post hoc tests (Tukey’s or Dunnett’s). Two-way ANOVA was performed for experiments involving two independent variables, such as genotype and treatment conditions. Outlier samples were identified and excluded from the analysis by the ROUT method (Q = 1%). The number of biological replicates (*n*) varied between experiments and is specified in the corresponding figure legends. A *p*-value < 0.05 was considered statistically significant.

## RESULTS

### Aldosterone/MR signaling activation induces primary VICs differentiation, inflammation, fibrosis and osteogenesis

We first aimed to evaluate the contribution of MR signaling to key pathophysiological processes involved in AS, including inflammation, VIC activation, fibrosis, and osteogenesis, using primary VICs isolated from rat AVs. Treatment with aldosterone induced a time-dependent upregulation of genes associated with VIC-activation and fibrosis genes, such as *Acta2*, *Vim*, and *Col1a1*. Transcriptional response was observed at 24 hours, which had further increased after six days of continuous exposure. These effects were abolished by cotreatment with finerenone, a selective non-steroidal (ns)-mineralocorticoid receptor antagonist (MRA), confirming that the observed changes in gene expression were mediated by MR activation (Figure 1A).

**Figure 1.**
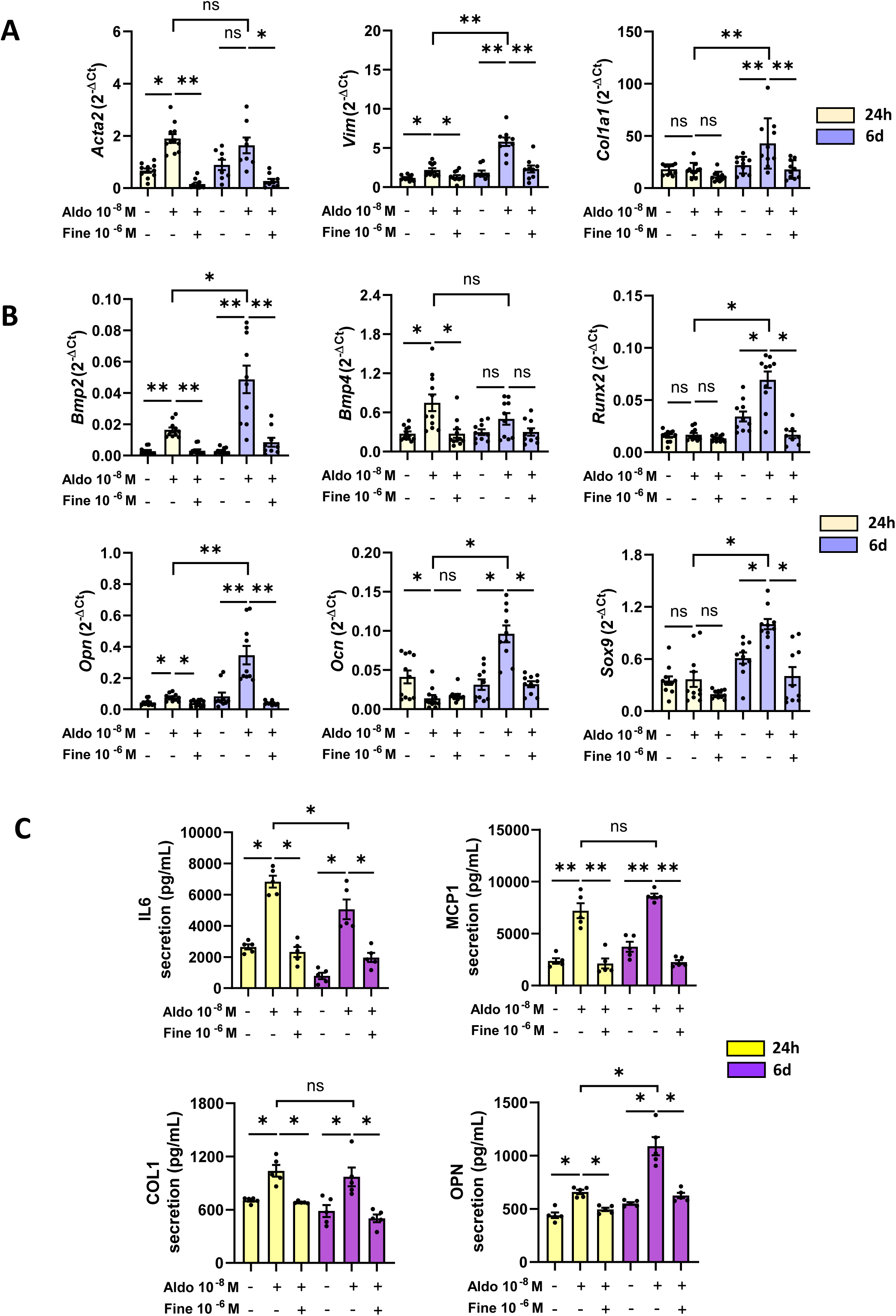
Effects of Aldo/MR signaling on AS pathological markers. mRNA expression (2^−ΔCt^, normalized to *Gapdh*) and protein secretion levels in VICs after 24 h or 6 days of treatment with Aldo (10⁻⁸ M) ± Fine (10⁻⁶ M). **A**, Relative expression of differentiation and matrix extracellular component (MEC) remodeling markers: *Acta2* (Actin Alpha 2, Smooth Muscle), *Vim* (Vimentin), *Col1a1* (Collagen type I alpha 1). **B**, Relative expression of early osteogenic markers: *Bmp2* (Bone Morphogenetic Protein 2), *Bmp4* (Bone Morphogenetic Protein 4), *Runx2* (Runt-related Transcription Factor 2), *Opn* (Osteopontin); and late osteogenic and chondrogenic markers: *Ocn* (Osteocalcin), and *Sox9* (SRY-box Transcription Factor 9), respectively. **C,** Protein secretion measured in cell supernatants: IL6, MCP1/CCL2, COL1, and OPN. Data are presented as mean ± SEM (n = 5–8 biological replicates). Statistical analysis was performed using one-way ANOVA followed by Dunnett’s post hoc test. *p < 0.05, **p < 0.01, ***p < 0.001.

Concomitant treatment with aldosterone for 24 hours triggered an early upregulation of the osteogenic markers *Bmp2*, *Bmp4* and *Opn*. Continuous exposure for six days led to a further increase in Bmp2 and Opn expression and a marked induction of *Runx2*, whereas no further change in Bmp4 expression was observed. These effects were significantly attenuated by finerenone (Figure 1B). By contrast, the expression of markers of late-stage osteogenesis, such as *Ocn*, and of the chondrogenic marker *Sox9*, increased only after prolonged stimulation with aldosterone. This delayed response was also abolished by the ns-MRA (Figure 1B). The role of inflammatory cytokines in the initiation of AS is well established. Our findings demonstrate that challenging VIC with aldosterone induces an inflammatory response by increasing not only the expression (Figure S1A) but also the secretion of proinflammatory cytokines, such as IL6 and MCP1/CCL2, together with structural and osteogenic proteins, such as COL1 and OPN (Figure 1C). These changes were also dependent on MR signaling, underscoring the critical role of MR in regulating these processes.

### Serum from CKD rats induces primary VICs differentiation, inflammation, fibrosis and osteogenesis via the MR signaling

We further investigated the contribution of CKD to AS progression by using pooled serum samples from sham-operated and CKD rats to treat healthy primary rat VICs. Mean aldosterone levels were much higher in pooled serum from CKD rats (567.5 pg/mL) than in pooled serum from sham-operated controls (55.5 pg/mL).

Exposure of VICs to CKD rat serum for six days led to an increase in the expression of VIC-activation and fibrotic markers, including Acta2, Vim, and Col1a1 (Figure 2A), and upregulation of early osteogenic genes *Bmp2*, *Bmp4*, *Runx2* and *Opn*, late-stage osteogenic markers, such as *Ocn*, and the chondrogenic marker *Sox9* (Figure 2B).

**Figure 2.**
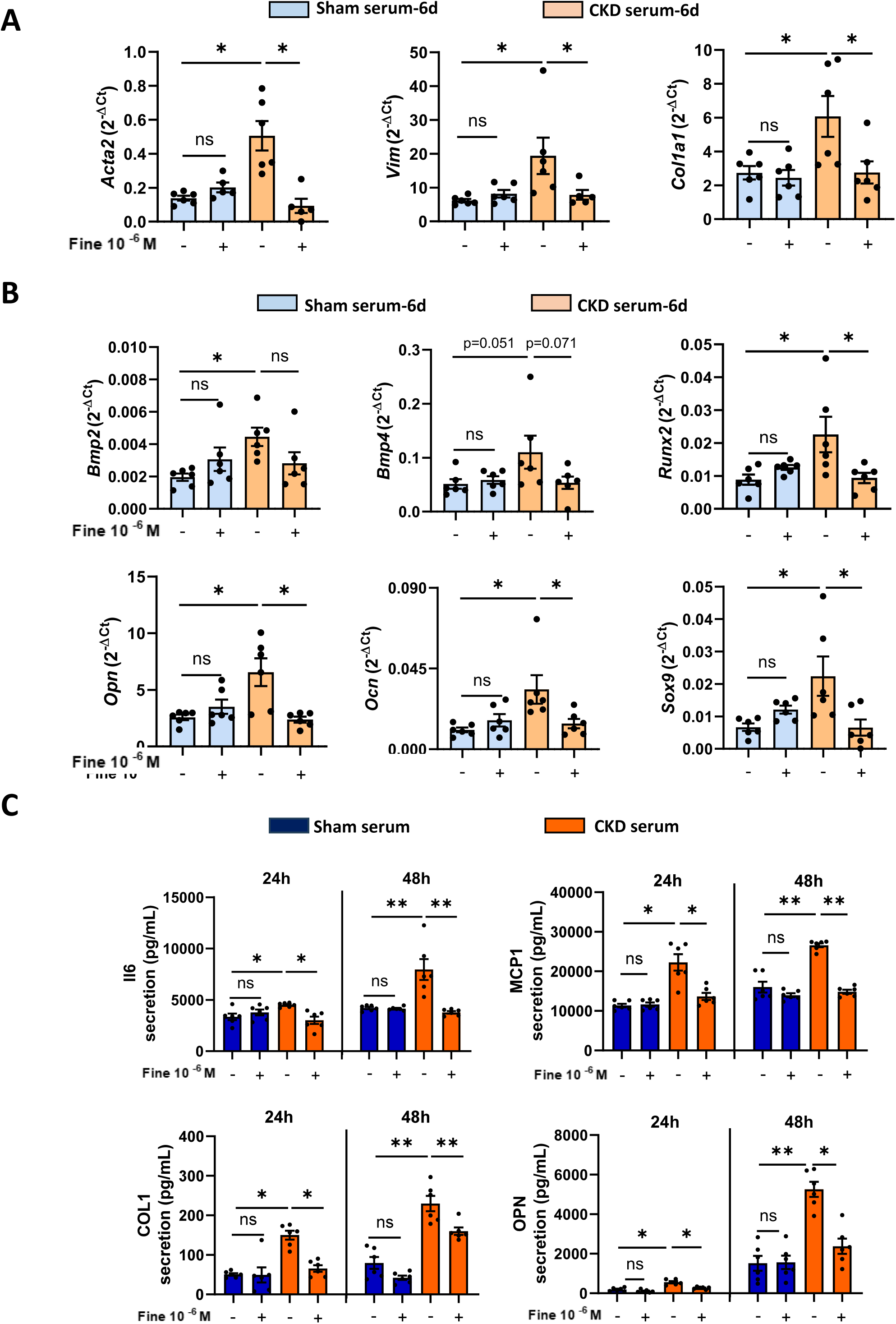
CKD serum effects on AS pathological markers. VICs exposed to CKD serum for the indicated times. **A-B,** Relative mRNA expression of EMC markers; *Acta2, Vim, Col1a1* and osteogenic*; Bmp2, Bmp4, Runx2; Opn, Sox9, Ocn* after 6 days of VICs treatment. Expression levels were normalized to *Gapdh.* **C,** IL6, CCL2/MCP1, COL1 and OPN secretion. Data are shown as mean ± SEM, n = 6 biological replicates. Statistical analysis was performed using two-way ANOVA followed by Sidak’s post hoc test. *p < 0.05, **p < 0.01, **p < 0.001.

Except for upregulation of *Bmp2*, all these effects were inhibited by the ns-MRA finerenone, demonstrating the involvement of MR signaling. By contrast, we observed no significant change in mRNA levels for inflammatory cytokines *Il6* and *Ccl2* after six days of treatment (Figure S1B). Protein secretion analysis revealed a marked increase in IL6, MCP1/CCL2, COL1, and OPN levels, this response peaking at 48 hours. These effects were fully abolished by finerenone, confirming their dependence on MR activation (Figure 2C).

### CKD induces differentiation, osteogenesis and fibrosis in the aortic valve via MR-signaling

We analyzed AVs from CKD rats, with and without treatment with the MRA eplerenone (Eple), to translate our findings for the rat primary VIC *in vitro* model to an *in vivo* setting. *In vivo* studies confirmed renal failure, demonstrated by increases in plasma levels of urea (Figure S1C). Immunohistochemical (IHQ) analysis (Figure 3A) revealed an increase in expression of VIC activation markers, such as αSMA/ACTA2, and high levels of osteogenic markers, including BMP2, BMP4 and RUNX2, and of the chondrogenic marker SOX9 in CKD rats. These alterations were significantly attenuated by eplerenone treatment. In addition, fibrosis, as assessed by Sirius Red staining (SR), increased markedly in CKD rats, in an MR-dependent manner. We used polarized light microscopy to quantify thick and thin collagen fibers involved in fibrotic remodeling. We found that numbers of both types of fibers were high in CKD rat AVs and were decreased by MRA treatment (Figure 3B).

**Figure 3.**
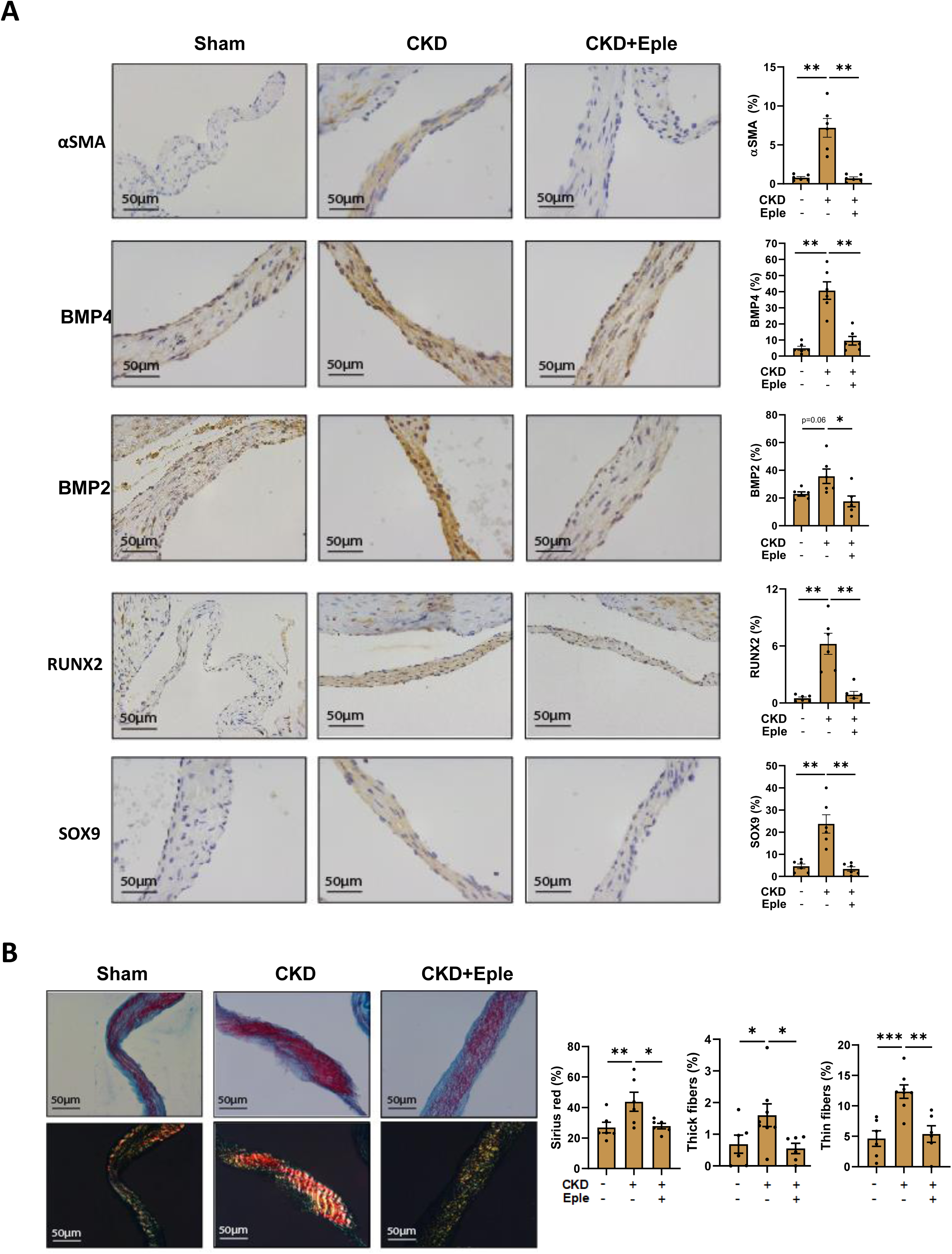
AV osteogenesis and fibrosis in CKD rats. AVs obtained from CKD and Sham rats generated as described in methods section. Representative microphotograph at 40× magnification of one leaflet of the rat AV stained with **(A)** DAB-based immunohistochemistry and (**B)** Sirius Red (SR) staining to assess fibrosis, visualized under both brightfield and polarized light to distinguish thick and thin collagen fibers. Quantification is presented as mean ± SEM, showing the percentage of positively stained area normalized to the total leaflet surface. n = 6 biological replicates. Statistical analysis was performed using one-way ANOVA followed by Dunnett’s post hoc test. *p < 0.05, **p < 0.01, **p < 0.001.

### NGAL is upregulated by MR activation in rat AVs and VICs and has osteogenic and inflammatory effects

Given the role of NGAL as an MR target, we then investigated the regulation of NGAL in response to aldosterone treatment. Activation of MR signaling by aldosterone in rat primary VICs induced a time-dependent increase in both expression and secretion of NGAL after 24 h or 6 days of stimulation (Figure 4A). VICs treated with pooled serum samples from CKD rats also displayed higher levels of NGAL gene expression and protein secretion than those treated with sham-operated rat serum (Figure 4B). These effects were abolished by the MRA finerenone, confirming that NGAL acts downstream from MR signaling (Figure 4A-B). IHQ analysis in the *in vivo* model demonstrated significantly stronger NGAL staining in AVs of CKD rats, which was attenuated by eplerenone treatment (Figure 4C and S1E).

**Figure 4.**
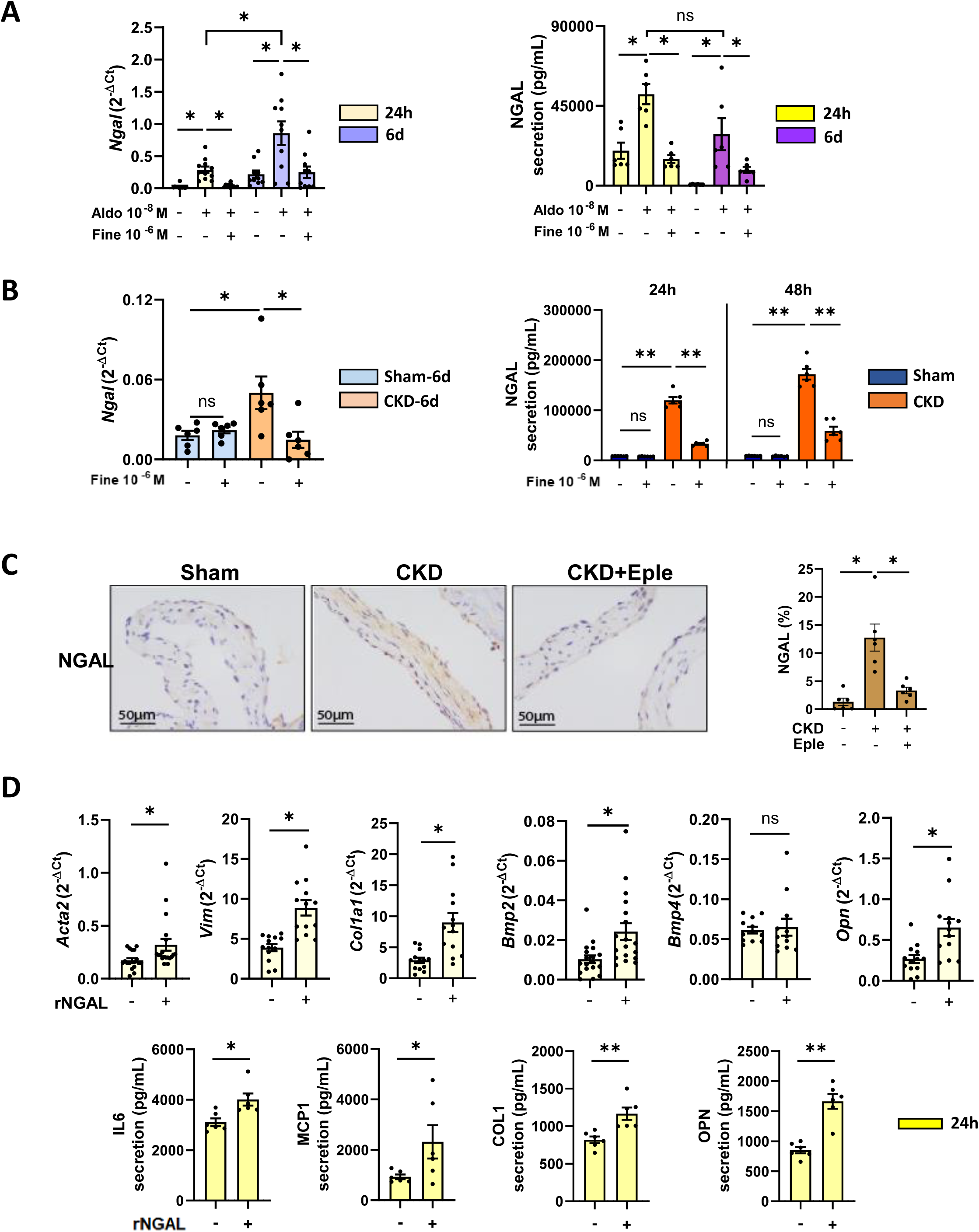
The presence of the MR-related target NGAL in Vics and AVs. NGAL expression/secretion in VICs treated with **(A)** Aldo (10^-8^ M) ± Fine (10^-6^ M) for 24h or 6 days and **(B)** CKD serum for 6 days, 24h or 48h, respectively. **C,** NGAL IHC in Sham or CKD rat AV ± Eple. **D,** Effect of recombinant (r)NGAL on mRNA (*Acta2, Vim, Col1a1, Bmp2, Bmp4, Opn*) and cytokine secretion after 24h-treatment. mRNA levels were normalized to *Gapdh* housekeeping gene. Data are shown as mean ± SEM, n = 6 - 12 biological replicates. Statistical analysis was performed using one-way ANOVA followed by Dunnett’s post hoc test or Student’s t-test, as appropriate. *p < 0.05, **p < 0.01, **p < 0.001.

We next explored the role of NGAL in VICs by treating cells with recombinant rat NGAL (rNGAL) for 24 hours. This treatment significantly increased expression of VIC activation markers (*Acta2*, *Vim*) (Figure 4D), a fibrotic marker (*Col1a1*), osteogenic markers (*Bmp2*, *Bmp4*, *Opn*, *Ocn*) (Figure 4D and S1F), and inflammatory cytokines, including MCP1/CCL2 and IL6, at both mRNA and secreted protein levels (Figure S1G and 4D). Secretion levels were also high for structural proteins, such as COL1 and the osteogenic OPN (Figure 4D). These results highlight the complex, multifaceted role of NGAL in modulating processes relevant to the pathophysiology of AS.

### NGAL mediates the effects of MR on differentiation, osteogenesis and inflammation in VICs and AVs

We further assessed the role of NGAL in mediating MR-induced effects in VICs, using primary VICs isolated from NGAL-knockout (KO-NGAL) rats. RT-qPCR and ELISA analyses confirmed complete absence of NGAL expression and secretion in KO-NGAL VICs (Figure S2A). After 24 hours of stimulation with aldosterone, KO-NGAL VICs displayed a reduced induction of expression of VIC activation markers (*Acta2*, *Vim*) and early osteogenic markers (*Bmp4*, *Opn*), whereas Bmp2 expression remained unaffected (Figure 5A). These effects were maintained up to six days of aldosterone challenge (Figure S2B). Following prolonged exposure to aldosterone over six days, KO-NGAL VICs also displayed attenuated expression of late-stage fibrotic (*Col1a1*), osteogenic (*Runx2*, *Ocn*), and chondrogenic (*Sox9*) markers relative to WT VICs (Figure 5B and 2C). Furthermore, analysis of secreted proteins after 24 hours of aldosterone treatment revealed a decrease in release of inflammatory cytokines, such as IL6 and MCP1/CCL2, and in levels of fibrotic and osteogenic markers, such as COL1 and OPN, in KO-NGAL cells (Figure 5C). These results were also reflected at gene expression level (Figure S2D). These results support the role of NGAL as a key mediator of MR-driven pathological remodeling in VICs.

**Figure 5.**
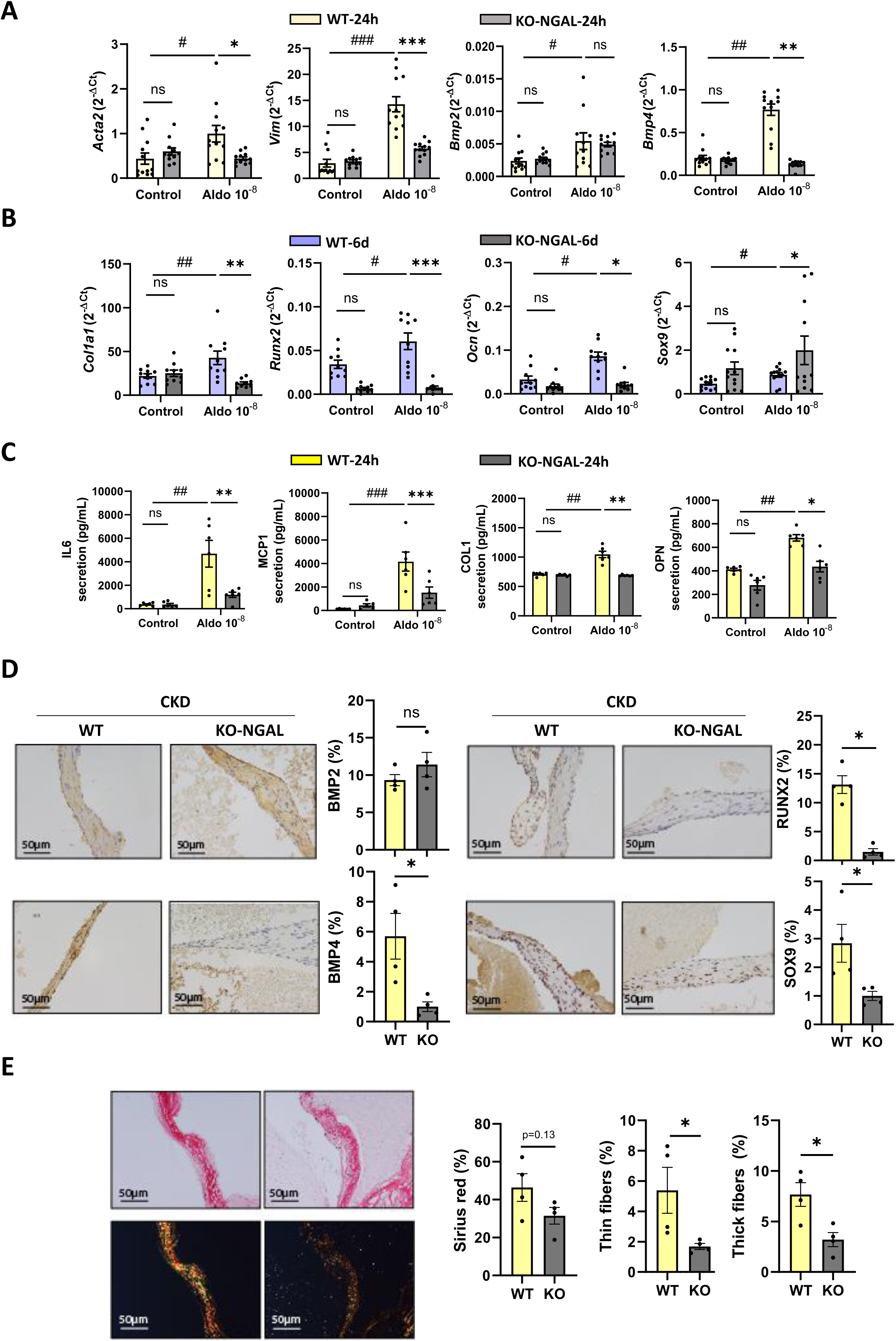
The role of NGAL in the pathological AS-related processes induced by MR signaling. **A-B,** Gene expression normalized to *Gapdh* in Wild-type (WT) and NGAL-knockout (KO-NGAL) VICs treated with Aldo (10⁻⁸ M) for 24h/6d as indicated, and **(C)** protein secretion at 24h. Representative microphotograph at 40× magnification of one leaflet of the rat AV with **(D)** DAB-based immunohistochemistry and **(E)** Sirius red (SR) staining to assess fibrosis, visualized under both brightfield and polarized light to distinguish thick and thin collagen fibers. Data are shown as mean ± SEM, n = 6–12 biological replicates. Statistical analysis was performed using two-way ANOVA followed by Sidak’s post hoc test or Student’s t-test, as appropriate. *p < 0.05, **p < 0.01, ***p < 0.001.

We extended these findings *in vivo*, by evaluating the impact of NGAL deletion on AV remodeling in CKD rats. Immunohistochemical analyses mirrored *in vitro* observations in VICs: levels of protein for osteogenic markers BMP4, RUNX2, and SOX9 were lower in AVs of KO-NGAL CKD rats than in WT-CKD rats, whereas BMP2 expression was unaffected (Figure 5D and S2E). Sirius Red staining further demonstrated that AV fibrosis levels were lower in NGAL-deficient animals, with markedly lower numbers of both thick and thin collagen fibers in AVs of KO-NGAL CKD rats than in those of WT-CKD rats (Figure 5E and S2E). Collectively, these findings support a central role for NGAL in promoting osteogenic differentiation and fibrotic remodeling in CKD, while highlighting its potential as a therapeutic target for prevention of pathological valve calcification.

### The TLR4-MyD88 innate immunity pathway is activated in response to aldosterone-induced MR signaling in rat AVs and VICs

Given that inflammation plays a key role in early stages of AS development and the changes in cytokine levels observed in response to aldosterone treatment, we investigated the potential involvement of specific inflammatory pathways downstream from MR activation. Based on previous Nanostring® analyses in VICs targeting fibrotic and inflammatory genes conducted by our team, we focused on the TLR4–MyD88 pathway. RT-qPCR analysis revealed upregulation of *Tlr4*, *Myd88*, *Irak1* and *Irak4* in response to MR activation by aldosterone (Figure S3A). Components of the MAP2K1 pathway were also upregulated (Figure S3A). This upregulation was blunted by MR inhibition with the ns-MRA finerenone (Figure S3A).

WB analysis further confirmed the increase in TLR4 and MyD88 protein levels following aldosterone treatment (Figure 6A). VICs displayed a trend toward enhanced NFκB phosphorylation, along with elevated levels of phosphorylated (p)-ERK, a key effector of the MAPK pathway (Figure 6A). We further characterized NFκB pathway activation by assessing levels of mRNA for its subunits. *Rela* (p65) and *Nfkb1* (p50) expressions were significantly upregulated at 24 hours, all effects being blunted by treatment with finerenone (Figure S3A).

**Figure 6.**
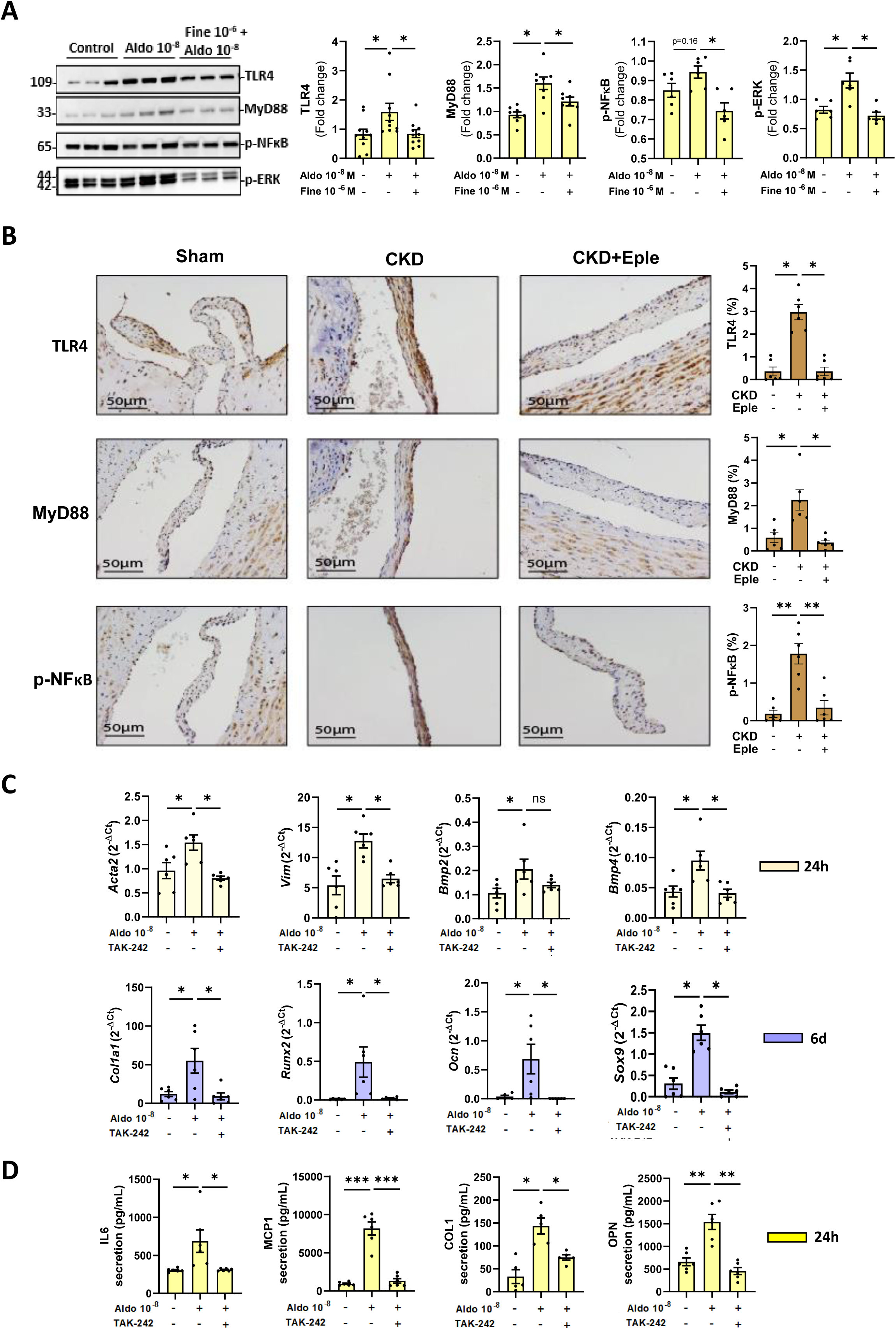
Identification of inflammatory pathways induced by Aldo/MR signaling. **A,** WB analysis of inflammatory pathways in VIC treated or not with Aldo (10^-8^ M) ± Fine (10^-6^ M) for 24h: TLR4 (Toll-like receptor 4), MyD88 (Myeloid Differentiation Primary Response 88), p-NFκB (Nuclear Factor kappa-light-chain-enhancer of activated B cells), p-ERK (Extracellular signal-Regulated Kinase). **B,** IHC of TLR4, MyD88 and p-NFκB in the AV of CKD male rats. **C-D,** mRNA levels or protein secretion of differentiation (*Acta2, Vim,* COL1), osteogenic (*Bmp2, Bmp4, Runx2, Ocn, Sox9, OPN*) and inflammatory (IL6, MCP1) markers at different time-points (yellow; 24h, purple; 6d) in the presence of Aldo (10^-8^ M) ± TAK-242 (TLR4 inhibitor, 1 µM). Data were normalized to *Gapdh*. Data are shown as mean ± SEM, n = 6 biological replicates. Statistical analysis was performed using one-way ANOVA followed by Tukey and Dunnett’s post hoc tests. *p < 0.05, **p < 0.01, ***p < 0.001.

The involvement of the TLR4–MyD88 and MAPK pathways was further supported by upregulation of the genes listed above in VICs treated with serum from CKD rats, an effect blunted by MR blockade (Figure S3B). Finally, immunohistochemical analysis of AVs from CKD rats corroborated these *in vitro* findings, showing MR-dependent increases in levels of TLR4, MyD88, and phosphorylated (p)-NFκB proteins (Figure 6B).

To extend our findings, we investigated the involvement of the TLR4 innate immune pathway in mediating aldosterone/MR signaling effects, using TAK-242, a selective TLR4 inhibitor. WB and RT-qPCR analyses confirmed effective inhibition of TLR4 signaling and its downstream targets (MyD88, p-ERK, p-NFκB) in presence of aldosterone and TAK-242 (Figure S3D-E). TLR4 blockade also highlighted the contribution of TLR4 signaling to aldosterone-induced changes in levels of differentiation (*Acta2*, *Vim*), osteogenic (*Bmp4*, *Runx2*, *Ocn*), fibrotic (*Col1a1*), and chondrogenic markers (*Sox9*) (Figure 6C). TAK-242 also blunted the increased expression of inflammatory genes (*Il6*, *Ccl2*) induced by aldosterone (Figure S3E). Notably, expression of *Bmp2* and *Opn* remained unaffected by TLR4 inhibition at mRNA level (Figure 6C and S3E). Finally, increased secretion of inflammatory cytokines (IL6, MCP1/CCL2) and of COL1 and OPN by aldosterone was regulated by TLR4 activation downstream from the MR, as demonstrated by the inhibitory effect of TAK-242 (Figure 6D).

### Aldosterone/MR signaling in VICs is mediated by the NGAL-dependent activation of the TLR4-MYD88 axis

As *Ngal* gene inactivation and TLR4 signaling affected aldosterone-induced responses, we investigated a possible link between NGAL and the TLR4 signaling pathway. We treated VICs with recombinant NGAL (rNGAL) for 24 hours. This treatment increased protein and mRNA levels of Tlr4 and Myd88 (Figure S4A) and led to activation of downstream targets (Figure S4B), supporting a role for NGAL in modulating TLR4 signaling. In addition, NGAL levels measured in presence of aldosterone and TAK-242 indicated that NGAL levels were unchanged by TLR4 inhibition, suggesting NGAL probably acts upstream from TLR4 signaling (Figure S4C).

We further investigated the role of NGAL by analyzing impact of aldosterone-stimulation in KO-NGAL VICs. Contrary to results obtained for WT VICs, aldosterone induced no increase in levels of TLR4, MyD88, p-NFκB, and p-ERK proteins (Figure 7A). These results were supported by RT-qPCR analysis showing a decrease in expression of *Tlr4*, *Myd88*, and downstream effectors, including *Irak1*, *Irak4*, *Rela*, *Nfkb1* and *Map2k1* (Figure S4D).

**Figure 7.**
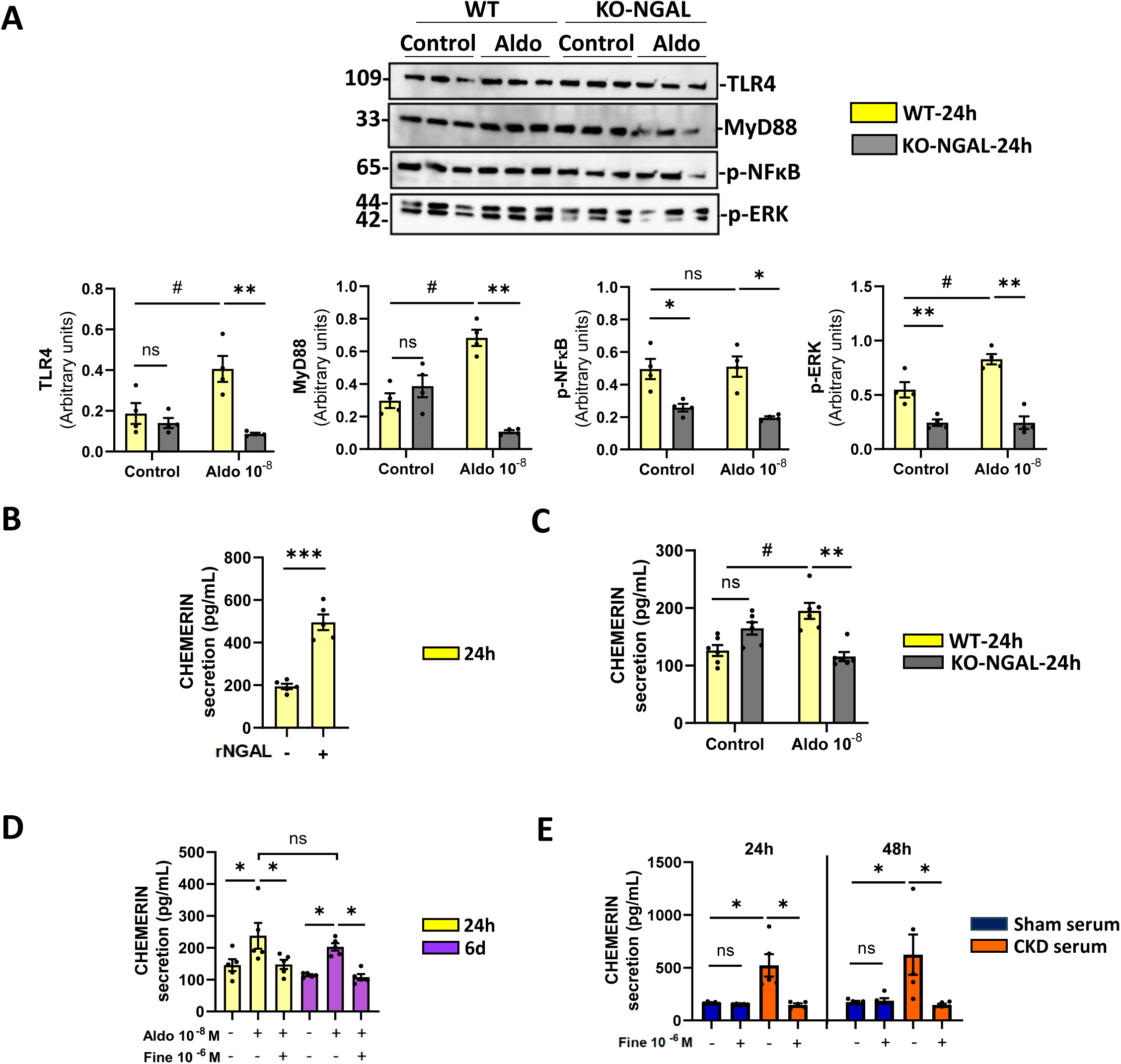
Identification of the potential effectors acting downstream the MR signaling. **A,** Wild-type (WT) and NGAL-knockout (KO-NGAL) VICs were exposed to Aldo (10⁻⁸ M) for 24 h, followed by protein analysis. Chemerin secretion after treatment of VICs with rNGAL **(B)**, or comparison of WT and KO-NGAL VICs exposed to aldosterone for 24 hours **(C).** Chemerin secretion evaluated under two conditions: **(D)** VICs treated with aldosterone for 24 hours or 6 days or **(E)** VICs exposed to serum from CKD rats for the indicated durations. Data are shown as mean ± SEM, n = 4-10 biological replicates. Statistical analysis was performed using one-way ANOVA followed by Dunnett’s post hoc test, two-way ANOVA followed by Sidak’s post hoc test or Student’s t-test, as appropriate. *p < 0.05, **p < 0.01, ***p < 0.001.

NGAL has not been identified as a direct ligand for TLR4, but previous studies have linked it to production of chemerin, a molecule identified as a potential endogenous TLR4 activator. We therefore assessed chemerin production in response to rNGAL treatment. RT-qPCR analysis revealed an increase in chemerin (*Rarres2*) expression (Figure S4E) and ELISA confirmed an increase in secretion (Figure 7B). Supporting these findings, exposure of KO-NGAL VICs to aldosterone demonstrated a requirement of NGAL for aldosterone/MR-induced chemerin (*Rarres2*) expression and secretion (Figure S4F and 7C).

We therefore assessed chemerin levels in response to MR signaling. RT-qPCR analysis revealed an increase in chemerin (*Rarres2*) expression (Figure S4G-H) and ELISA confirmed an increase in secretion (Figure 7D-E) of this molecule following treatment with aldosterone and CKD serum. These effects were blocked by the ns-MRA finerenone (Figure 7D-E and S4G-H).

Together, these results underscore the crucial role of NGAL in activating the TLR4 pathway in response to aldosterone, contributing to inflammation, remodeling, and osteogenesis in VICs.

### Chemerin signaling induces the TLR4 pathway in response to aldosterone/MR activation

As chemerin has been identified as a potential endogenous TLR4 activator that could mediate its effect through its specific receptor Chemokine-Like Receptor 1 (CMKLR1), we studied the role of recombinant chemerin (rChem) in the regulation of the innate immunity pathway. Results exhibited a significant upregulation of TLR4 and MYD88 at both the mRNA and protein levels (Figure 8A). This upregulation was accompanied by an increase in the expression of additional effectors acting downstream from TLR4, including *Rela* and *Nfkb1* (Figure 5A). Recombinant chemerin also enhanced the expression of markers of differentiation and fibrosis (*Acta2, Vim, Col1a1*), osteogenic markers (*Bmp2, Bmp4*), and inflammatory cytokines (*Ccl2, Il6*). These effects were dependent on TLR4 signaling (Figure S5B-C).

**Figure 8.**
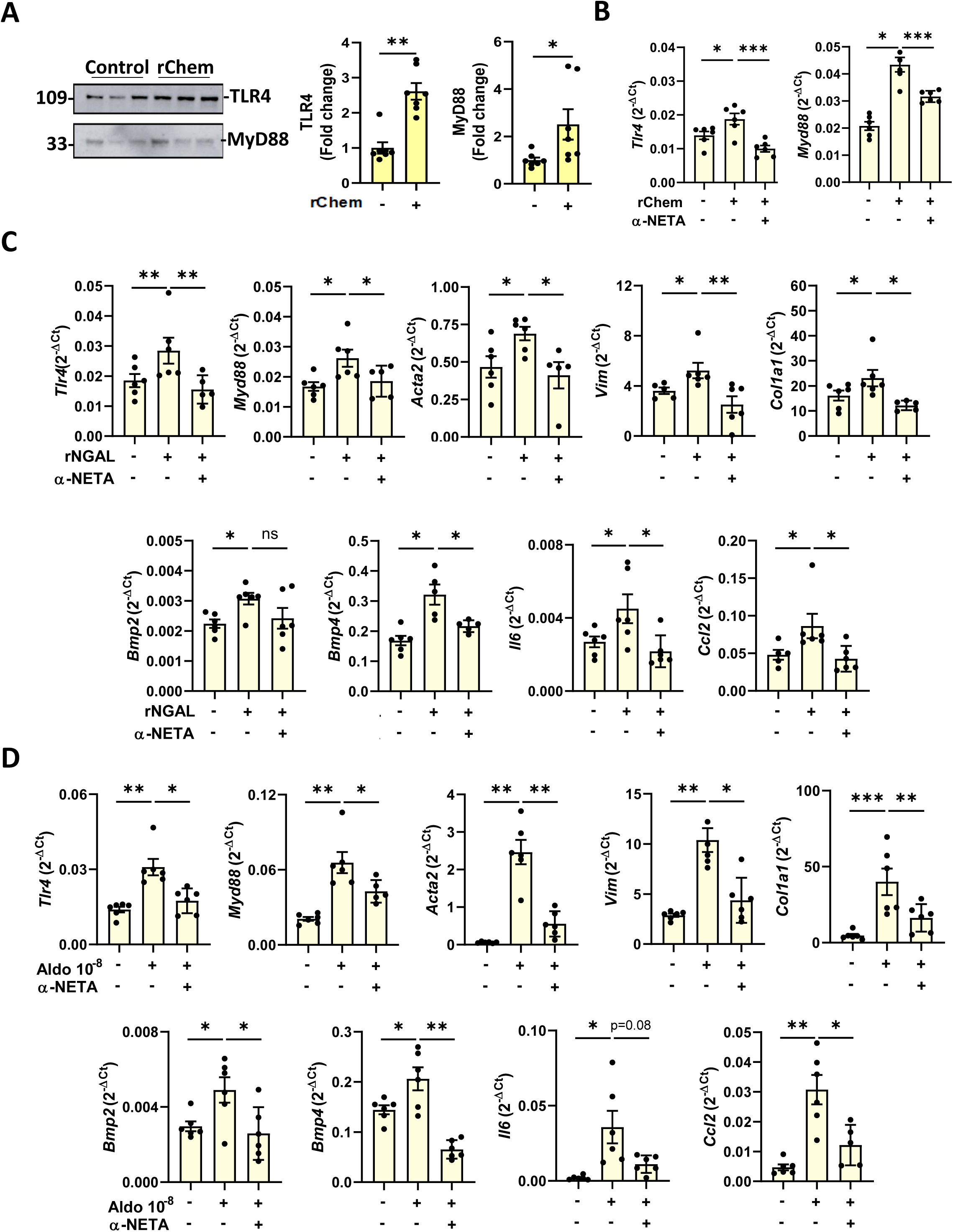
Role of chemerin on TLR4 signalling in aldosterone-induced fibrosis, osteogenesis and inflammation. VICs were treated for 24 h with recombinant chemerin (rChemerin, 50 nM) ± α-Neta (10 µM), an antagonist of the chemerin receptor CMKLR1. **A,** WB analyses of TLR4 and MyD88 protein. **B,** Gene expression levels of *Tlr4* and *Myd88*. VICs were also exposed to recombinant (r)NGAL (500 ng/mL) ± α-Neta (10 µM) for 24 h, and the expression of **(C)** *Tlr4* and *Myd88*, along with (genes involved in fibrotic differentiation (*Acta2*, *Vim*, *Col1a1*) and markers of osteogenesis (*Bmp2*, *Bmp4*) and inflammation (*Il6*, *Ccl2*), was evaluated. In parallel, VICs were stimulated with aldosterone (Aldo, 10^-8^ M) ± α-Neta (10 µM) for 24 h, followed by assessment of **(D)** *Tlr4*, *Myd88*, fibrotic, osteogenic, and inflammatory gene expression. Data are presented as mean ± SEM, with n = 4–7 biological replicates. Statistical significance was determined by one-way ANOVA with Tukey’s post hoc test or Student’s *t*-test, as appropriate. *p* < 0.05, p < 0.01, *p < 0.001.

Furthermore, blockade of the chemerin receptor (CMKLR1) using the inhibitor α-NETA demonstrated that the effects of recombinant chemerin (rChem) on *Tlr4*, *Myd88*, and their downstream signaling pathways are mediated specifically through CMKLR1 (Figure 8B and S5D). RT-qPCR analysis confirmed that inhibition of the chemerin receptor with α-NETA effectively abolished rChem-induced alterations in VIC differentiation, osteogenesis, and inflammatory responses (Figure S5D-E). Moreover, gene expression profiling of VICs treated with rNGAL in the presence of the CMKLR1 inhibitor further underscored the critical role of the chemerin receptor in upregulating *Tlr4*, *Myd88*, and their downstream targets (Figure 8C and S5F), as well as in driving VIC differentiation, fibrosis, osteogenesis, and inflammation downstream of MR signaling (Figure 8C). Finally, the CMKLR1 inhibitor also negated the effects induced by aldosterone on *Tlr4*, *Myd88*, and associated downstream effectors (Figure 8D and S5G), along with the consequent activation of VICs, fibrosis, osteogenesis, and inflammation (Figure 8D).

Together, these results reveal a complex interplay between NGAL and chemerin in activating the innate immune TLR4 signaling pathway in response to aldosterone/MR activation, contributing to inflammation, remodeling, and osteogenesis in VICs.

### Expression of the MR/NGAL/chemerin/TLR4 pathway is enhanced in a CKD-dependent manner in human valves

RT-qPCR analysis of AV tissues from patients undergoing valve replacement surgery and at different stages of CKD revealed a stage-dependent upregulation of the MR (*NR3C2*), together with expression of the *TLR4* and *MYD88* genes (Figure 9A). Complementary ELISA studies demonstrated a progressive increase in the levels of the NGAL and chemerin proteins (Figure 9A). Furthermore, *in vitro* analysis revealed that aldosterone treatment increased *TLR4* and *MYD88* gene expression in human VICs in an MR-dependent manner (Figure 9B), thereby supporting the findings obtained in the animal model.

**Figure 9.**
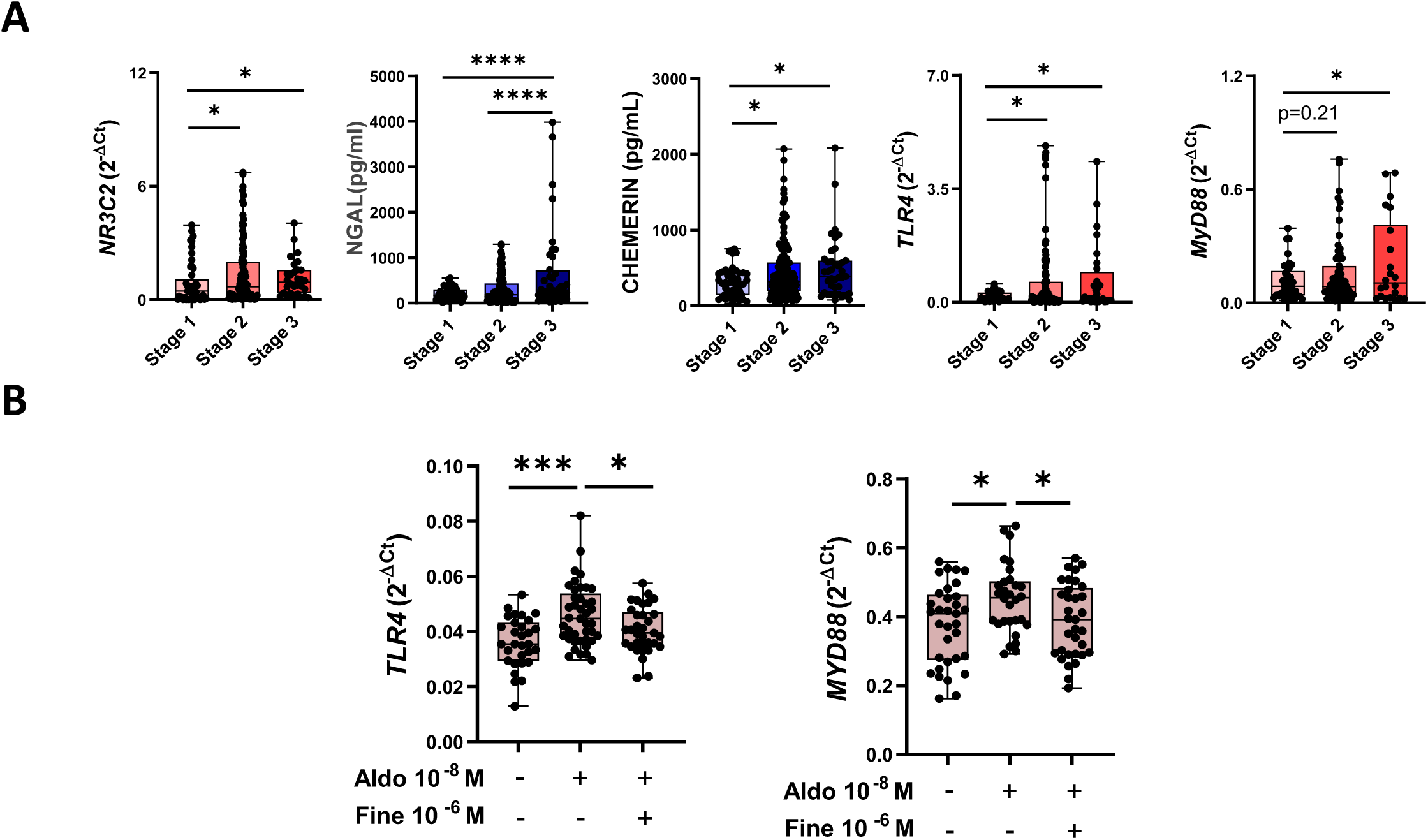
The NGAL/Chem/TLR4 pathway in human AV and VICs. **A,** Protein and mRNA expression measured by ELISA or qPCR in human AV, gene expression normalized to *GAPDH, B-ACTIN and HPRT.* **B,** mRNA levels of *TLR4* and *MYD88* in human VICs treated with Aldo 10^-8^ M ± Fine 10^-6^ M for 24h. Data are shown as mean ± SEM, n = 4-10 biological replicates. Statistical analysis was performed using one-way ANOVA followed by Tukey’s post hoc test. *p < 0.05, **p < 0.01, ***p < 0.001.

## DISCUSSION

This study identifies MR signaling as a novel and potentially actionable mechanism in the pathogenesis of AS, particularly in the setting of CKD. Our *in vitro* data demonstrate that aldosterone, as well as CKD serum with elevated aldosterone levels, promotes VIC activation, differentiation, inflammation, and osteogenic responses in a time- and MR-dependent manner. These findings are further supported by our *in vivo* CKD rat model, underscoring the pathophysiological relevance of MR signaling *in vivo*. Notably, these results suggest that pharmacological blockade of the MR with MRA may represent a promising therapeutic approach to mitigate AS progression in patients with concomitant CKD. Although clinical data specifically addressing AS are currently lacking, the beneficial cardiovascular and renal outcomes observed with MRA in large clinical trials such as FIGARO-DKD and FIDELIO-DKD, conducted in patients with CKD and type 2 diabetes, provide indirect but compelling support for the potential clinical translation of our findings^21,22^.

Our findings are consistent with previous studies establishing the aldosterone/MR axis as a key mediator of vascular inflammation, oxidative stress, and calcification, key processes implicated in the onset and progression of cardiovascular disease^9,10^. Recent evidence has extended the role of the MR pathway to valvular disease, indicating that it may contribute not only to vascular damage but also to valvular remodeling. In stenotic human AVs, MR is expressed in VICs, where its activation triggers cell-specific pro-osteogenic and pro-inflammatory responses. These effects are modulated by factors such as sex and metabolic comorbidities, including diabetes mellitus^12,19^. In agreement with these studies, our data indicate that MR expression is elevated in human AVs at early stages of CKD (stage 2–3), further supporting the notion that MR signaling contributes to valvular pathology in this patient population. MR blockade with the non-steroidal MRA finerenone has been shown to exert protective vascular effects in a type 1 diabetes CKD model using Wistar rats, including the downregulation of osteogenic, fibrotic, inflammatory, and oxidative stress pathways in renal tissue, as well as in perivascular and perirenal adipose tissue^21,22^.

Our findings reveal a temporally dynamic response to aldosterone in VICs. Early exposure elicits a rapid inflammatory response, consistent with the established paradigm in which immune cell infiltration and the release of cytokines, such as IL-1β, IL-6, TNF-α, and IFNs, trigger VIC activation and early osteogenic differentiation during AS^23,24,25^. Our data indicate that this initial phase is MR-dependent and precedes the more advanced osteogenic programming seen upon prolonged aldosterone stimulation, consistent with the late-stage features of AS. *Ex vivo* experiments have further validated the role of MR signaling in AV calcification in the context of CKD, a condition traditionally associated with passive mineral deposition due to a dysregulation of phosphate and calcium homeostasis^6,13^. Importantly, our data suggest that MR activation is an active, cell-mediated component of the process of calcification, probably constituting a mechanistic link between CKD and valvular disease.

Taken together, these findings highlight the importance of MR signaling as a key mediator of VIC pathobiology and as a newly identified contributor to the AS progression associated to CKD. This study adds to the growing body of evidence supporting MR antagonism as a promising strategy for attenuating the progression of AS, especially in high-risk populations. Indeed, the therapeutic relevance of this approach is further supported by recent clinical trials, such as FIDELIO and FIGARO, demonstrating cardiovascular and renal protective effects of the ns-MRA finerenone in patients with type 2 diabetes and CKD^21,22^. However, the impact of MRAs on valvular remodeling has yet to be assessed in this population.

This study identifies NGAL as a key downstream effector of MR signaling in the pathogenesis of AS, both *in vitro* and *in vivo*, in the AVs of CKD rats. Our data demonstrate that NGAL contributes to VIC differentiation, fibrosis, calcification, and inflammation, positioning it as a critical mediator of MR-driven valvular remodeling. NGAL was initially identified in neutrophils, but is now known to be expressed in diverse cell types, including epithelial and renal tubular cells, hepatocytes, cardiomyocytes, endothelial cells, vascular smooth muscle cells, fibroblasts, macrophages, and dendritic cells^26^. In particular, NGAL has been identified as a transcriptional target of MR signaling in various organs, including the gut, kidney, and heart^17,27,28^. NGAL expression has also been reported in porcine and human VICs under hypoxic conditions^15^ and, more recently, in stenotic human AV tissue^17^. NGAL expression is strongly associated with pro-inflammatory, fibrotic, and osteogenic molecular profiles in cardiovascular diseases^29^. Consistent with this observation, Jover et al.^18^ demonstrated a positive correlation between NGAL levels and inflammatory, oxidative stress, osteogenic, and calcific burden in human AS specimens. In addition, high circulating NGAL levels have been associated not only with disease progression, but also with cardiac remodeling and high levels of fibrotic marker expression in patients with aortic stenosis^30^. In the context of cardiorenal syndrome, a positive correlation has been found between plasma NGAL levels in non-diabetic patients undergoing hemodialysis and osteogenesis and calcification of the aortic root, and between these NGAL levels and serum calcium levels^31^.

In this study, we extend these findings by showing that recombinant NGAL (rNGAL) mimics aldosterone-induced effects in VICs at early time points, suggesting that NGAL mediates part of the downstream response to MR activation. Previous studies reported differential effects between intracellular and extracellular NGAL^32^, but our observations are consistent with those of Martínez-Martínez et al., who showed that rhNGAL triggers inflammatory and profibrotic responses in human cardiac fibroblasts^17^. Similarly, Jover et al.^18^ observed that rhNGAL increased the expression of *CCL2* and *OPN* in human VICs cultured in calcifying conditions, although it did not upregulate additional inflammatory or osteogenic markers, probably due to the overriding effects of high inorganic phosphate levels in the culture medium^18^. These results suggest that NGAL may amplify existing calcific stimuli rather than initiating the full osteogenic program on their own. Importantly, our findings indicate that NGAL does not mediate the transcriptional upregulation of *Bmp2* in response to aldosterone, raising the possibility that *Bmp2* activation is either directly regulated by MR or involves alternative signaling crosstalk. This observation points to a more complex regulatory network, in which MR signaling orchestrates different transcriptional programs via NGAL-dependent and NGAL-independent pathways.

In summary, the data presented here identify NGAL as a crucial mediator of MR-driven valvular disease contributing to VIC activation, inflammation, fibrosis, and calcification. Given its upstream regulation by MR and its capacity to modulate key pathological processes in AS, NGAL is a promising biomarker and potential therapeutic target for MR-driven valve disease, particularly in patients with CKD.

In addition to elucidating the roles of MR signaling and its downstream effector NGAL, this study identifies activation of the TLR4–MyD88–pERK innate immune signaling pathway as a novel axis driving cell differentiation, inflammation, and osteogenesis during AS progression upon aldosterone/MR activation, particularly in the context of CKD. Innate immune signaling was first implicated in valvular disease in 2008 when an increase in the expression of TLR2 and TLR4 in stenotic AVs was reported^2^. The dysregulation of these receptors was subsequently linked to osteogenesis and congenital abnormalities, such as bicuspid aortic valve syndrome^3,33^. Multiple *in vitro* studies have since further elucidated the role of TLRs, particularly TLR4 and TLR3, in the pathophysiology of VICs. These receptors have been shown to drive VIC differentiation and calcification via canonical signaling pathways, including the NFκB and ERK pathways, and to be involved in crosstalk with additional pro-inflammatory cascades, such as the JAK–STAT and WNT pathways^24,25,34,35^. More recently, TLR4 and TLR3 have been implicated in mediating the metabolic reprogramming of VICs through their downstream effectors NFκB and ERK, and, in cooperation with JAK–STAT signaling, in supporting the energetic and biosynthetic demands of osteogenic and calcific processes during AS progression^36^. TLR4 has been implicated in aldosterone-induced injury, particularly in fibrotic and inflammatory responses in heart and kidney tissues^37^. Notably, the aldosterone-MR–TLR4 interaction has been shown to mediate fibrotic signaling through the production of proteoglycans that serve as endogenous TLR4 ligands^37,38,39^. Our data extend these findings to VICs by showing that aldosterone-MR–NGAL signaling activates TLR4–MyD88–pERK without any significant detectable activation of NFκB via phosphorylation. Previous studies also reported delayed NFκB phosphorylation or an absence of such phosphorylation following TLR4 stimulation^39^, but it is important to note that the transcriptional activity of NFκB is not strictly dependent on its phosphorylation status^40^. Consistent with this, we observed an increase in *Il6* mRNA levels in response to aldosterone, a well-established target of NFκB, suggesting functional activation of this transcriptional pathway despite the lack of detectable phosphorylation.

Our data indicate that NGAL acts upstream of TLR4 signaling, specifically through the induction of chemerin production, which in turn engages its receptor to activate TLR4 signaling. This intermediate step suggests a mechanistic pathway in which NGAL promotes TLR4-mediated responses via chemerin-dependent signaling. Previous studies have demonstrated NGAL induction downstream from TLR4 in models of renal injury^41^, but our findings suggest that upstream activation occurs, with MR-driven NGAL promoting TLR4–MyD88 activation. This conclusion is supported by published findings indicating that NGAL can stimulate innate immune responses via TLR4, primarily through iron dysregulation and oxidative stress^42,43^. A deeper analysis of the mechanisms of action of NGAL revealed that chemerin was secreted in response to MR–NGAL activation. Chemerin, an adipokine and chemoattractant protein involved primarily in the regulation of inflammation, metabolism and immune responses, has been identified as a direct activator of TLR4 capable of stimulating TLR4–NFκB signaling and cytokine expression in human synovial fibroblasts from patients with arthritis^44^. In the context of AS, Goñi-Olóriz et al.^19^ recently identified chemerin as a key mediator of aldosterone-induced inflammation, differentiation, and calcification in human VICs^19^.

Taken together, these results support a model in which MR activation leads to an increase in NGAL expression, in turn promoting chemerin production, leading to the activation of TLR4– MyD88 signaling. This cascade ultimately drives key pathological processes in AS, including inflammation, VIC differentiation, fibrosis, and osteogenesis. The identification of this signaling axis adds a new layer to our mechanistic understanding of valve calcification and identifies new potential targets for therapeutic intervention. Moreover, our data support a potential therapeutic benefit of MR blockade, limiting AV stenosis, in CKD patients. In addition, our findings suggest that targeting NGAL or blocking the chemerin receptor could also represent effective strategies to attenuate disease progression and its associated pathological features.

## Author Contributions

T.S.B, F.J and N.L.A provided the concept and design of research

T.S.B, M.S, M.G, and A.F.C performed experiments and prepared figures,

T.S.B, M.S, M.G, A.F.C, F.J and N.L.A analyzed data and interpreted results of experiments.

T.S.B, F.J and N.L.A drafted the manuscript.

F.J and N.L.A approved the final version of the manuscript.

## Acknowledgement and funding

This work was funded by grants from the Institut National de la Santé et de la Recherche Médicale (INSERM), Fondation pour la Recherche Médicale, ANR NGAL-HT (ANR-19-CE14-0013), ANR MIRAVALVE (ANR-23-CE14-0016-01), Instituto de Salud Carlos III, the European Regional Development Fund (ERDF) – “A way of making Europe” (PI24/01020), and the IRP INSERM Miravac-CKD grant. We also thank our collaborators for their work as well as team members, and unit colleagues for their valuable support throughout this project.

## Supplemental information

Supplemental methods:Tables S1-S5

Supplemental figures: Figure S1-S5 Uncropped gels

## Data Availability Statement

The data generated during and/or analyzed during the current study are available from the corresponding author upon reasonable request.

## List of nonstandard abbreviations and acronyms

ACTA2 / α-SMA: Actin Alpha 2, Smooth Muscle / Alpha-Smooth Muscle Actin
Aldo: Aldosterone
AS: Aortic Stenosis
AV: Aortic Valve
BMP2: Bone Morphogenetic Protein 2
BMP4: Bone Morphogenetic Protein 4
CCL2/MCP1: C-C Motif Chemokine Ligand 2 / Monocyte Chemoattractant Protein 1
CKD: Chronic Kidney Disease
CMKLR1: Chemokine-like receptor 1
COL1A1/COL1: Collagen Type I Alpha 1 Chain/ Collagen Type I
Eple: Eplenerone
ERK: Extracellular Signal-Regulated Kinase
Fine: Finerenone
IL6: Interleukin 6
IRAK1: Interleukin-1 Receptor-Associated Kinase 1
IRAK4: Interleukin-1 Receptor-Associated Kinase 4
MAPK: Mitogen-Activated Protein Kinase
MR: Mineralocorticoid Receptor
MRA: Mineralocorticoid Receptor Antagonist
MYD88: Myeloid Differentiation Primary Response 88
NFKB: Nuclear Factor Kappa-Light-Chain-Enhancer of Activated B Cells
NGAL: Neutrophil Gelatinase-Associated Lipocalin (also known as Lipocalin 2 / LCN2)
rNGAL: Recombinant Neutrophil Gelatinase-Associated Lipocalin
rChem: Recombinant Chemerin
OCN: Osteocalcin
OPN: Osteopontin
RELA: v-rel Avian Reticuloendotheliosis Viral Oncogene Homolog A (p65 subunit of NF-κB)
RUNX2: Runt-Related Transcription Factor 2
SOX9: SRY-Box Transcription Factor 9
TAK-242: Toll-Like Receptor 4 Inhibitor (also known as Resatorvid)
TLR4: Toll-Like Receptor 4
VIC: Valvular Interstitial Cell
VIM: Vimentin

## REFERENCES

1. Vahanian, A., Beyersdorf, F., Praz, F., Milojevic, M., Baldus, S., Bauersachs, J., & Wokakowski, W. ESC/EACTS Guidelines for the management of valvular heart disease: Developed by the Task Force for the management of valvular heart disease of the European Society of Cardiology (ESC) and the European Association for Cardio-Thoracic Surgery (EACTS). Eur. Heart J. 12;43(7):561–632 (2022).

2. Meng, X., Ao, L., Song, Y., Babu, A., Yang, X., Wang, M., & Fullerton, D. A. Expression of functional Toll-like receptors 2 and 4 in human aortic valve interstitial cells: potential roles in aortic valve inflammation and stenosis. Am. J. Physiol. Cell Physiol. 294(1): 29–35 (2008).

3. Yang X., Fullerton D.A., Su X., Ao L., Cleveland J.C. & Meng X. Pro-osteogenic phenotype of human aortic valve interstitial cells is associated with higher levels of Toll-like receptors 2 and 4 and enhanced expression of bone morphogenetic protein 2. J. Am. Coll. Cardiol. 53(6):491–500 (2009).

4. Alushi, B., Curini, L., Christopher, M. R., Grubitzch, H., Landmesser, U., Amedei, A., & Lauten, A. Calcific Aortic Valve Disease-Natural History and Future Therapeutic Strategies. Front. Pharmacol. 11:685 (2020).

5. Ternacle J., Côté N., Krapf L., Nguyen A., Clavel M.A., Pibarot P.. Chronic Kidney Disease and the Pathophysiology of Valvular Heart Disease. Can. J. Cardiol. 35(9):1195–207 (2019).

6. Rattazzi M., Bertacco E., Del Vecchio A., Puato M., Faggin E., Pauletto P.. Aortic valve calcification in chronic kidney disease. Nephrol. Dial. Transplant. 28(12):2968–76 (2013).

7. Verma A., Vaidya A., Subudhi S., Waikar S.S. Aldosterone in chronic kidney disease and renal outcomes. Eur. Heart J. 43(38):3781–91 (2022).

8. Aros C., & Remuzzi G. The renin-angiotensin system in progression, remission and regression of chronic nephropathies. J. Hypertens. 20(3):S45–53 (2002).

9. Jaffe I.Z., Tintut Y., Newfell B.G., Demer L.L., Mendelsohn M.E. Mineralocorticoid receptor activation promotes vascular cell calcification. Arterioscler. Thromb. Vasc. Biol. 27(4):799–805 (2007).

10. Young M. J. Mechanisms of mineralocorticoid receptor-mediated cardiac fibrosis and vascular inflammation. Curr. Opin. Nephrol. Hypertens. 17(2):174–80 (2008).

11. Gkizas, S., Koumoundourou, D., Sirinian, X., Rokidi, S., Mavrilas, D., Koutsoukos, P., Papalois, A., Apostolakis, E., Alexopoulos, D., & Papadaki, H. Aldosterone receptor blockade inhibits degenerative processes in the early stage of calcific aortic stenosis. Eur. J. Pharmacol. 642(1):107–12 (2010).

12. Matilla L. et al. Sex-Related Signaling of Aldosterone/Mineralocorticoid Receptor Pathway in Calcific Aortic Stenosis. Hypertens. 79(8):1724–37 (2022).

13. Linefsky JP., O’Brien K.D., Katz R., Boer I.H., Barasch E., Jenny N.S., siscovick D.D. & Kestenbaum B. Association of Serum Phosphate Levels with Aortic Valve Sclerosis and Annular Calcification: the Cardiovascular Health Study. J. Am. Coll. Cardiol. 58(3):291–7 (2011).

14. Bolignano, D., Lacquaniti, A., Coppolino, G., Donato, V., Campo, S., Fazio, M. R., Nicocia, G., & Buemi, M. Neutrophil gelatinase-associated lipocalin (NGAL) and progression of chronic kidney disease. Clin. J. Am. Soc. Nephrol. 4(2):337–44 (2009).

15. Bonnard, B., Ibarrola, J., Lima-Posada, I., Fernández-Celis, A., Durand, M., Genty, M., Lopez-Andrés, N., & Jaisser, F. Neutrophil Gelatinase-Associated Lipocalin From Macrophages Plays a Critical Role in Renal Fibrosis Via the CCL5 (Chemokine Ligand 5)-Th2 Cells-IL4 (Interleukin 4) Pathway. Hypertens. 79(2):352–64 (2022).

16. Swaminathan G, Krishnamurthy VK, Sridhar S, Robson DC, Ning Y, Grande-Allen KJ. Hypoxia Stimulates Synthesis of Neutrophil Gelatinase-Associated Lipocalin in Aortic Valve Disease. Front. Cardiovasc. Med. 6:156 (2019).

17. Martinez-Martinez, E., Ibarrola, J., Calvier, L., Fernandez-Celis, A., Leroy, C., Cachofeiro, V., Rossignol, P., & Lopez-Andres, N. Aldosterone Target NGAL (Neutrophil Gelatinase-Associated Lipocalin) Is Involved in Cardiac Remodeling After Myocardial Infarction Through NFκB Pathway. Hypertens. 70(6):1148–56 (2017).

18. Jover E. et al. Sex-dependent expression of neutrophil gelatinase-associated lipocalin in aortic stenosis. Biol. Sex. Differ. 13(1):71 (2022).

19. Goñi-Olóriz M. et al. Chemerin is a new sex-specific target in aortic stenosis concomitant with diabetes regulated by the aldosterone/mineralocorticoid receptor axis. Am. J. Physiol. Heart. Circ Physiol. 328(3):H639–47 (2025).

20. Sádaba, J. R., Martinez-Martinez, E., Arrieta, V., Álvarez, V., & Fernández-Celis, A. Role for Galectin-3 in Calcific Aortic Valve Stenosis. J. Am. Heart Assoc. 5(11):e004360 (2016).

21. Sanz-Gómez M. et al. Finerenone protects against progression of kidney and cardiovascular damage in a model of type 1 diabetes through modulation of proinflammatory and osteogenic factors. Biomed. Pharmacother. 168:115661 (2023).

22. Bakris GL. et al. Effect of Finerenone on Chronic Kidney Disease Outcomes in Type 2 Diabetes. N. Engl. J. Med. 383(23):2219–29 (2020).

23. Raddatz M.A., Madhur M.S., & Merryman W.D. Adaptive immune cells in calcific aortic valve disease. Am. J. Physiol Heart. Circ. Physiol. 317(1):H141–55 (2019).

24. Parra-Izquierdo, I., Sánchez-Bayuela, T., Castaños-Mollor, I., López, J., Gómez, C., San Román, J. A., Sánchez Crespo, M., & García-Rodríguez, C. Clinically used JAK inhibitor blunts dsRNA-induced inflammation and calcification in aortic valve interstitial cells. FEBS. J. 288(22):6528–42 (2021).

25. Parra-Izquierdo, I., Castaños-Mollor, I., López, J., Gómez, C., San Román, J. A., Sánchez Crespo, M., & García-Rodríguez, C. Lipopolysaccharide and interferon-γ team up to activate HIF-1α via STAT1 in normoxia and exhibit sex differences in human aortic valve interstitial cells. Biochim. Biophys. Acta Mol. Basis Dis. 1865(9):2168–79 (2019).

26. Romejko K., Markowska M., & Niemczyk S. The Review of Current Knowledge on Neutrophil Gelatinase-Associated Lipocalin (NGAL). Int. J. Mol. Sci. 24(13):10470 (2023).

27. Bonnard B. et al. Antifibrotic effect of novel neutrophil gelatinase-associated lipocalin inhibitors in cardiac and renal disease models. Sci. Rep. 11(1):2591 (2021).

28. Amamou A. et al. Mineralocorticoid receptor activation contributes to intestinal fibrosis through neutrophil gelatinase-associated lipocalin in preclinical models. Nat. Commun. 16:6318 (2025).

29. Ramos-Mozo P. et al. Increased plasma levels of NGAL, a marker of neutrophil activation, in patients with abdominal aortic aneurysm. Atheroscl. 220(2):552–6 (2012).

30. Bäz L. et al. Serum Biomarkers of Cardiovascular Remodelling Reflect Extra-Valvular Cardiac Damage in Patients with Severe Aortic Stenosis. Int. J. Mol. Sci. 21(11):4174 (2020).

31. Wei K. et al. Association of plasma neutrophil gelatinase-associated lipocalin and thoracic aorta calcification in maintenance hemodialysis patients with and without diabetes. BMC. Nephrol. 23(1):156 (2022).

32. Tong Z., Wu X., Ovcharenko D., Zhu J., Chen C.S., & Kehrer J.P. Neutrophil gelatinase-associated lipocalin as a survival factor. Biochem J. 391(Pt 2):441–8 (2005).

33. Balistreri C.R. et al. Deregulation of TLR4 signaling pathway characterizes Bicuspid Aortic valve syndrome. Sci. Rep. 9(1):11028 (2019).

34. Jarrett, M. J., Houk, A. K., McCuistion, P. E., Weyant, M. J., Reece, T. B., Meng, X., & Fullerton, D. A. Wnt Signaling Mediates Pro-Fibrogenic Activity in Human Aortic Valve Interstitial Cells. Ann. Thorac. Surg. 112(2):519–25 (2021).

35. Zhang Y., Peng W., Ao X., Dai H., Yuan L., Huang X., & Zhou Q. TAK-242, a Toll-Like Receptor 4 Antagonist, Protects against Aldosterone-Induced Cardiac and Renal Injury. PLoS One. 10;10(11):e0142456. (2015).

36. Sánchez-Bayuela T. et al. Inflammation via JAK-STAT/HIF-1α Drives Metabolic Changes in Pentose Phosphate Pathway and Glycolysis That Support Aortic Valve Cell Calcification. Arterioscler. Thromb. Vasc. Biol. 45(7):e232–49 (2025).

37. Palacios-Ramirez, R., Soulie, M., Fernandez-Celis, A., Nakamura, T., Boujardine, N., Bonnard, B., Bamberg, K., Lopez-Andres, N., & Jaisser, F. Mineralocorticoid receptor (MR) antagonist eplerenone and MR modulator balcinrenone prevent renal extracellular matrix remodeling and inflammation via the MR/proteoglycan/TLR4 pathway. Clin. Sci. Lond. 138(16):1025–38 (2024).

38. Nakamura T., Bonnard B., Palacios-Ramirez R., Fernández-Celis A., Jaisser F., López-Andrés N.. Biglycan Is a Novel Mineralocorticoid Receptor Target Involved in Aldosterone/Salt-Induced Glomerular Injury. Int. J. Mol. Sci. 23(12):6680 (2022).

39. Jarrett, M. J., Yao, Q., Venardos, N., Weyant, M. J., Reece, T. B., Meng, X., & Fullerton, D. A. Simvastatin down-regulates osteogenic response in cultured human aortic valve interstitial cells. J Thorac. Cardiovasc. Surg. 161(4):e261–e271. (2021).

40. Smith E.L., Somma D., Kerrigan D., Mclntyre Z., Cole J.J., Liang K.L., Kiely P.A., Keeshan K. & Carmody, R. C. The regulation of sequence specific NF-κB DNA binding and transcription by IKKβ phosphorylation of NF-κB p50 at serine 80. Nucleic Acids Res. 47(21):11151–63 (2019).

41. Wu C., Lv C., Chen F., Ma X., Shao Y., & Wang Q. The function of miR-199a-5p/Klotho regulating TLR4/NF-κB p65/NGAL pathways in rat mesangial cells cultured with high glucose and the mechanism. Mol. Cell. Endocrinol. 417:84–93 (2015).

42. Mori K. et al. Endocytic delivery of lipocalin-siderophore-iron complex rescues the kidney from ischemia-reperfusion injury. J. Clin. Invest. 115(3):610–21 (2005).

43. Cowland J.B., & Borregaard N. Molecular Characterization and Pattern of Tissue Expression of the Gene for Neutrophil Gelatinase-Associated Lipocalin from Humans. Genomics. 45(1):17–23 (1997).

44. Eisinger, K., Bauer, S., Schäffler, A., Walter, R., Neumann, E., Buechler, C., Müller-Ladner, U., & Frommer, K. W. Chemerin induces CCL2 and TLR4 in synovial fibroblasts of patients with rheumatoid arthritis and osteoarthritis. Exp. Mol. Pathol. 92(1):90–6 (2012).

